# Microglia mediate contact-independent neuronal pruning via secreted Neuraminidase-3 associated with extracellular vesicles

**DOI:** 10.1101/2023.08.21.554214

**Authors:** Corleone S. Delaveris, Catherine L. Wang, Nicholas M. Riley, Sherry Li, Rishikesh U. Kulkarni, Carolyn R. Bertozzi

## Abstract

Neurons communicate with each other through electrochemical transmission at synapses. Microglia, the resident immune cells of the central nervous system, can prune these synapses through a variety of contact-dependent and -independent means. Microglial secretion of active sialidase enzymes upon exposure to inflammatory stimuli is one unexplored mechanism of pruning. Recent work from our lab showed that treatment of neurons with bacterial sialidases disrupts neuronal network connectivity. Here, we find that activated microglia secrete Neuraminidase-3 (Neu3) associated with fusogenic extracellular vesicles. Furthermore, we show Neu3 mediates contact-independent pruning of neurons and subsequent disruption of neuronal networks through neuronal glycocalyx remodeling. We observe that *NEU3* is transcriptionally upregulated upon exposure to inflammatory stimuli, and that a genetic knock-out of *NEU3* abrogates the sialidase activity of inflammatory microglial secretions. Moreover, we demonstrate that Neu3 is associated with a subpopulation of extracellular vesicles, possibly exosomes, that are secreted by microglia upon inflammatory insult. Finally, we demonstrate that Neu3 is both necessary and sufficient to both desialylate neurons and decrease neuronal network connectivity. These results implicate Neu3 in remodeling of the glycocalyx leading to aberrant network-level activity of neurons, with implications in neuroinflammatory diseases such as Parkinson’s disease and Alzheimer’s disease.

**Graphical Abstract:** Neuroinflammation induces secretion of the sialidase Neu3 via extracellular vesicles from microglia that prune neuronal synapses and disrupt neuronal communication.

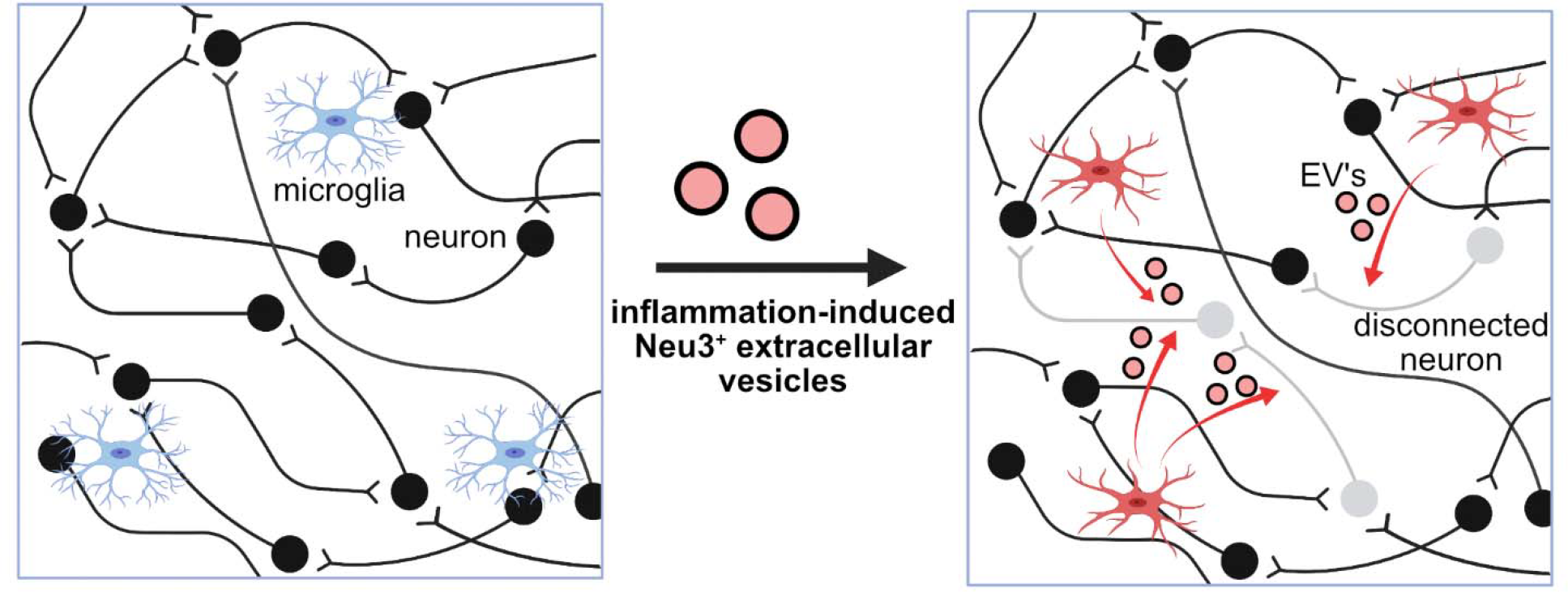

## Introduction

The brain is made of a vast interconnected and interdependent network of neurons, communicating information through synapses. Microglia, the resident immune phagocytes of the central nervous system, prune these synapses through several mechanisms, including direct and complement-mediated phagocytosis.^1–7^ These activities are upregulated in the context of neuroinflammatory pathologies, including Alzheimer’s disease and Parkinson’s disease.^8–10^ However, the specific mechanisms by which hyperinflammatory microglia mediate these effects remain unclear, especially in the context of how these actions impact neuronal networking and communication through synapses. Given that neurodegenerative diseases correlate with aberrant network-level neuronal activity,^9,10^ it is important to understand the molecular mechanisms by which inflammatory microglia regulate neuronal communication.

Neuroinflammation has been correlated with changes in the glycocalyx – the coating of sugars on cell surfaces – of both neurons and microglia.^11–15^ Sialic acids are a particular subset of bioactive sugars in the glycocalyx. They are known to modulate neuronal excitability and plasticity,^7,16^ and changes in sialylation state are associated with neuroinflammation and microglial activation.^16–22^ Upon exposure to inflammatory stimuli, microglia have been observed to release sialidase activity into the surrounding media, which effects desialylation,^17,22,23^ deposition of opsonizing factors,^18,23^ microglial activation,^16,23^ and phagocytosis of neurons.^17,23^ Additionally, our lab has recently identified sialylation state as a critical factor in maintaining neuronal excitability and network integration.^21^ Collectively, these observations point to the glycocalyx as a regulator of neuronal activity.

Herein, we tested the hypothesis that sialidases released by microglia could affect contact-independent neuronal pruning. We found that the peripheral membrane glycolipid sialidase Neuraminidase-3 (Neu3) is secreted by microglia upon activation by inflammatory stimuli. Neu3 was localized to a population of extracellular vesicles that are fusogenic with neurons. Using a voltage-sensing imaging dye, we found Neu3 is both necessary and sufficient to mediate neuronal pruning and the disconnection of neuronal networks. Based on these data, we propose a mechanism in which microglia secrete Neu3 to remodel neuronal glycocalyces to modulate neuronal connectivity. These results have implications for how neuroinflammation results in neuronal network dysfunction.

## Results and Discussion

### Activated microglia upregulate *NEU3* and require *NEU3* to desialylate neuronal glycocalyces

We and others have observed that microglial secretions possess sialidase activity and are capable of desialylating model cell lines^24^ and primary neurons (**Figure 1a,b**). Notably, these effects can be pharmacologically inhibited with zanamivir, which has inhibitory activity for human sialidases.^25^ Of the four mammalian sialidases, three have reported expression in the brain.^26^ To identify the sialidase(s) secreted by activated microglia responsible for this activity, we activated BV-2 murine microglia using lipopolysaccharide (LPS) and assessed relative mRNA expression of sialidase genes using qPCR. We observed a 50% increase in *NEU3* transcripts following activation (*p=*0.041) and statistically insignificant changes in *NEU1* and *NEU4* (*p*=0.11 and *p*=0.90, respectively) (**Figure 1c**). To investigate the role of the glycolipid sialidase Neu3^27^ in desialylating neurons, we generated *NEU3* (the gene encoding Neu3) knockout (KO) BV-2 microglia and compared the sialidase activity of wild-type (WT) and *NEU3* KO BV-2 secretions (**Figure S1**). *NEU3* KO conditioned media exhibited minimal sialidase activity compared to WT (*p*=0.043), as measured by Peanut Agglutinin (PNA) binding of terminal N-acetylgalactosamine exposed by desialylation on neuronal membranes (**Figure 1d**). Consistent with the observation that Neu3 is predominantly a glycolipid (e.g., ganglioside) sialidase,^27–30^ we did not observe significant changes in sialylated glycoproteins of neurons treated with conditioned media from WT versus *NEU3* KO microglia (**Table S1**).

**Figure 1.**
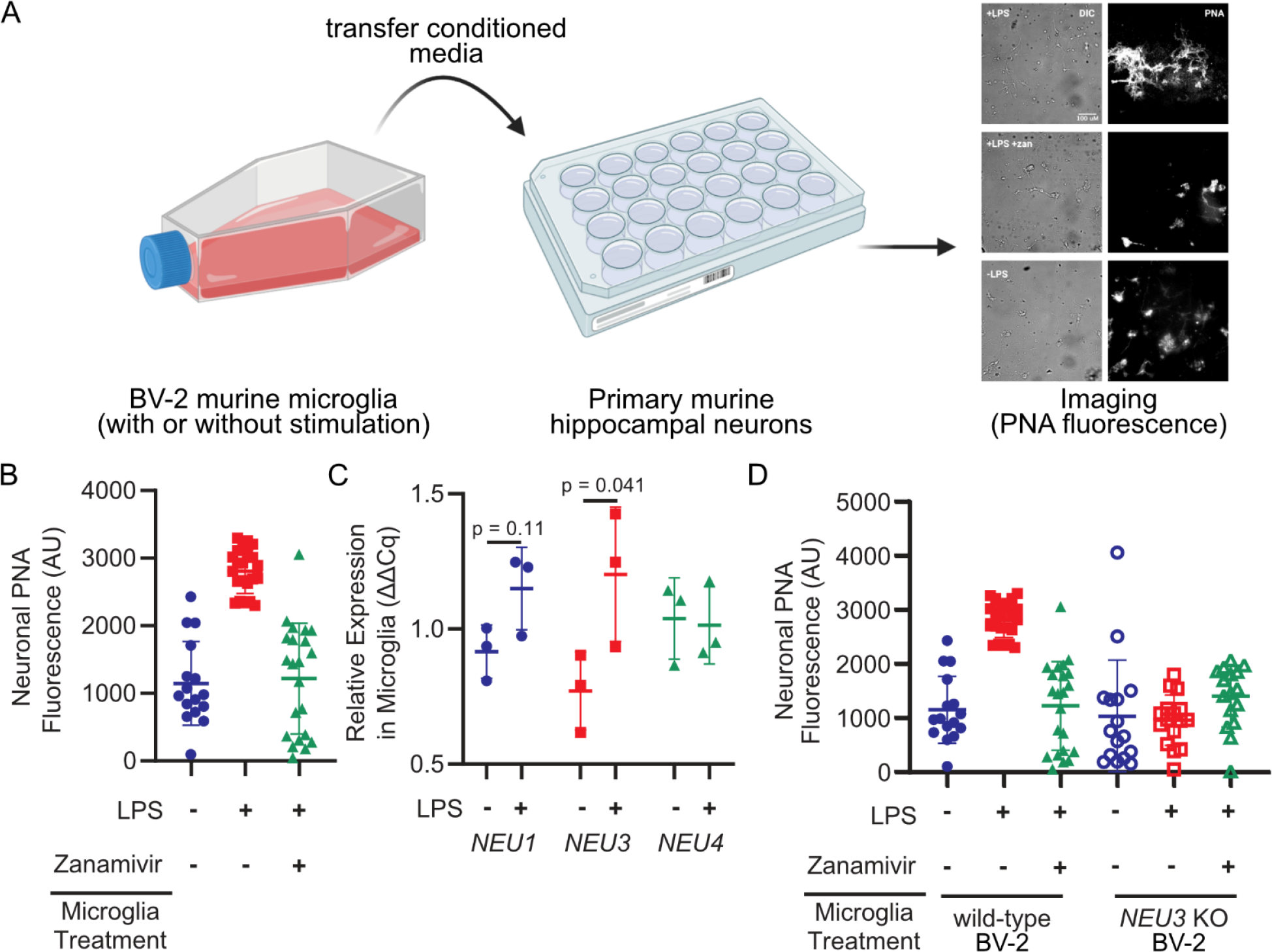
Microglia upregulate and NEU3 and active Neu3 is necessary for secreted sialidase activity. **(A, B)** Primary mouse hippocampal neurons were treated with conditioned media from resting or LPS-activated BV-2 microglia in the presence or absence of zanamivir. Representative scheme and images **(A)** and quantification of fluorescence **(B)** reveal that LPS-activation causes 3-fold increase in PNA signal compared to resting (+LPS vs. -LPS, *p*=0.045), an effect abrogated by pharmacological sialidase inhibition (-LPS vs. +LPS +zan, *p*=0.90; +LPS vs. +LPS +zan, *p*=0.042). Hypothesis testing performed with hierarchical permutation test, n=3 coverslips/condition, avg. 20 neurons/condition. **(C)** Quantification of transcript levels of *NEU1, NEU3*, and *NEU4* by qPCR in resting and LPS-activated BV-2 microglia (*NEU1, p*=0.11; *NEU3, p*=0.041; *NEU4, p*=0.90). **(D)** Neurons were treated with conditioned media from wild-type (WT) or *NEU3* knockout (*NEU3* KO) BV-2 microglia, with or without deoxy-2,3-anhydroneuraminic acid (DANA), and stained with peanut agglutinin (PNA). Media from activated WT microglia produced a 3-fold increase in desialylation compared to resting (- LPS vs. +LPS, *p*=0.043; +LPS vs. +LPS+zan, *p*=0.042) but media from *NEU3* KO microglia exhibited no significant change in desialylation in response to LPS or zanamivir (-LPS vs. +LPS, *p*=0.74; +LPS vs. +LPS+zan, *p*=0.12). n=3 coverslips/condition, 60 total WT cells, 48 total *NEU3* KO cells. Hypothesis tests performed with hierarchical permutation test.

As prior studies have implicated Neu1 translocation and desialylation in cis as a critical component of microglial activation,^14,15,23^ we sought to determine whether the loss of secreted sialidase activity in *NEU3* KO BV-2 cells was a consequence of impaired inflammatory activity. We observed that *NEU3* KO cells had impaired autodesialylation in response to LPS treatment compared to WT as measured by periodate labeling of sialic acids (p=0.51 and =0.038, respectively), but that both WT and *NEU3* KO cells upregulated TNF□to similar levels in (WT, *p*=0.002; *NEU3* KO, *p*=0.02) (**Figure S2**). Therefore, *NEU3* KO cells are still able to secrete inflammatory signals. TNF□secretion in both cell lines was inhibited by the pan-sialidase inhibitor deoxy-2,3-anhydroneuraminic acid (DANA), consistent with previous observations with Neu1.^14,23^ Given that the BV-2 microglia are still capable of secreting inflammatory molecules, the autodesialylation likely does not have a biological impact in this context. Moreover, these data implicate Neu3 as the secreted sialidase, rather than an upstream component.

### Microglial secreted Neu3 is associated with extracellular vesicles that fuse with neurons

Neu3 has been shown to behave as a peripheral membrane protein,^31^ with recent studies demonstrating that the enzyme is S-acylated.^32^ Additionally, microglia are known to secrete extracellular vesicles (EVs) upon activation.^33^ Therefore, we hypothesized that Neu3 might be secreted in association with EVs. To investigate this hypothesis, we isolated EVs from resting and activated microglia conditioned media using commercial lectin-based isolation kits. We confirmed that we were isolating EVs based on proteomics of the surfaceome, which identified known EV proteins, but we were unable to detect Neu3 directly (**Figure S3, Table S2**). To confirm that Neu3 colocalizes with EVs, we inserted a 3xFLAG-tag at on the endogenous NEU3 gene by homology-directed recombination (**Figure S4**). We then performed an immunocapture bead assay in which magnetic beads were functionalized with anti-murine CD63 to capture EVs and incubated with microglia-conditioned medium. Captured EVs were analyzed with anti-FLAG and either anti-CD9 or anti-CD81, both of which are used as EV markers.^34^ We observed a FLAG-positive population of EVs, the relative population of which significantly increased upon LPS-stimulation, suggesting that microglial Neu3 is secreted via EVs upon immune stimulus (**Figure 2a, S5**).

**Figure 2.**
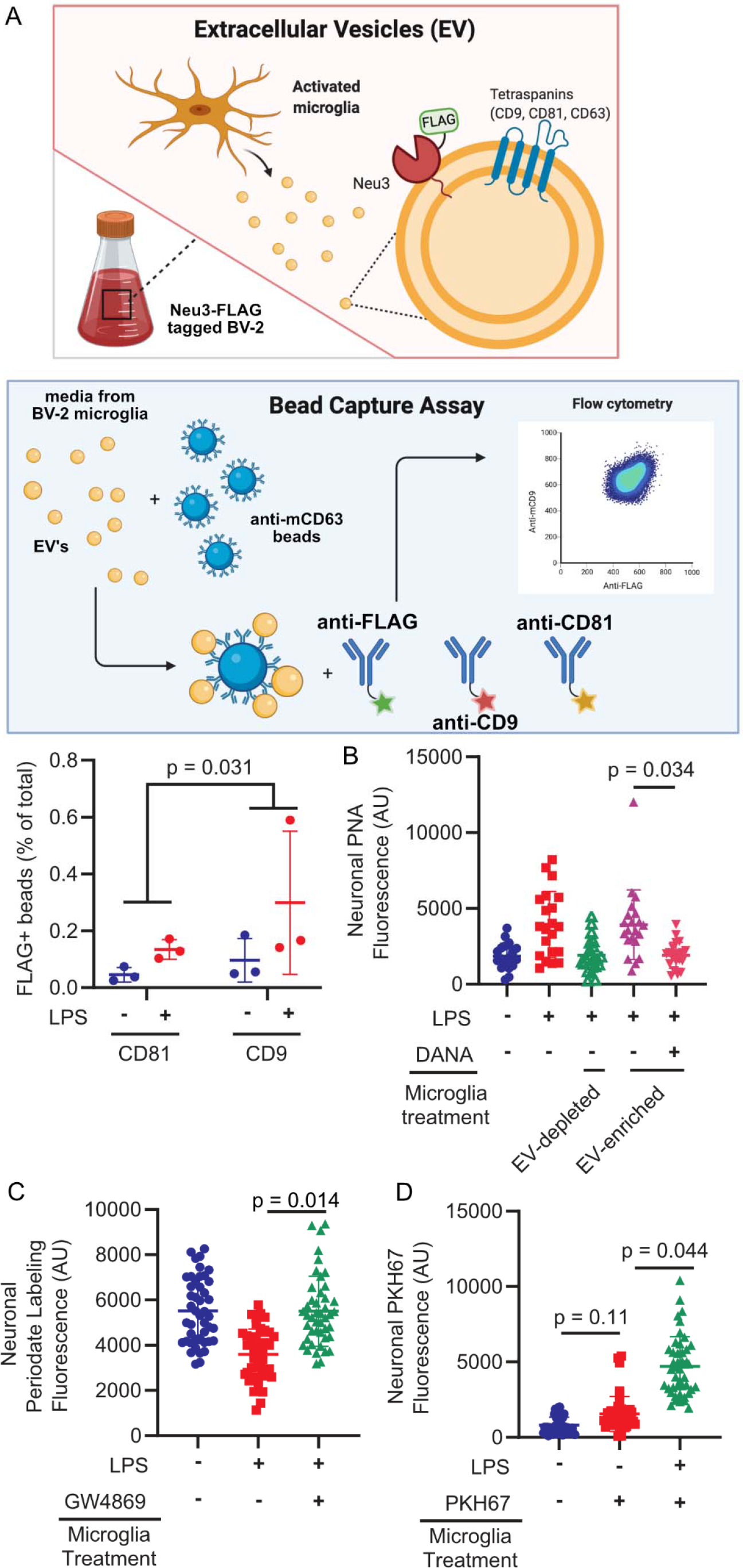
Neu3 is associated with microglia-derived fusogenic extracellular vesicles. **(A)** Endogenous NEU3 was FLAG-tagged in BV-2 microglia by homology-directed recombination. After exposure of BV-2 microglia with endogenously FLAG-tagged Neu3 to vehicle or LPS, EVs were captured on anti-mCD81 or anti-mCD9 coupled beads, labeled with fluorophore-coupled anti-FLAG or anti-mCD63, and analyzed by flow cytometry. Bead captured-EV’s demonstrate increased FLAG signal in LPS-treated microglia compared to resting microglia, indicating that *NEU3* colocalizes with EV markers and is released via EVs upon LPS-activation. (-LPS vs. +LPS, p-value = 0.031). n=3 wells/condition, 2 capture methods/well. **(B)** PNA staining of neuronal surfaces treated with EV-enriched or EV-depleted media of activated WT microglia reveals that EV-enriched fraction alone has sialidase activity (EV-enriched vs. EV-depleted, *p*=0.023; EV-enriched vs. EV-enriched+DANA, *p*=0.034). n=4 coverslips/condition, 7 cells/coverslip. **(C)** Periodate labeling of neuronal surface sialic acids reveals that pharmacologic inhibition of EV production with GW4869 abrogates sialidase activity of EV-enriched microglia media (-LPS vs. +LPS, *p*=0.027; -LPS vs. +LPS+GW4869, *p*=0.97; +LPS vs. +LPS+GW4869, *p*=0.014). n=3 coverslips/condition, 135 cells total. **(D)** Imaging of neurons treated with PKH67-stained microglial exosome demonstrates transferal of dye from EVs to neuronal membranes (Vehicle vs. -LPS, *p*=0.15; -LPS vs. +LPS, *p*=0.044). Hypothesis testing for all panels was performed using hierarchical permutation test. n=3 coverslips/condition, 15 cells/coverslip.

To confirm that EV-resident Neu3 is responsible for the sialidase activity on neurons, we treated neurons with EV-enriched or EV-depleted microglia conditioned media. We observed that EV-enriched fractions demonstrated significantly increased sialidase activity compared to EV-depleted fractions (*p*=0.023, **Figure 2b**), suggesting that EVs are responsible for desialylation. Consistent with this, pharmacological inhibition of extracellular vesicle production with GW4869^35^ decreased sialidase activity of the enriched fraction (*p*=0.014, **Figure 2c**). Furthermore, upon staining EVs with a membrane dye we observed robust dye transfer to neuronal membranes, indicating vesicle fusion with neurons (*p*=0.044, **Figure 2d**). These data suggest a model in which microglia expel extracellular vesicles containing Neu3, which fuse with neuronal membranes and cause desialylation of the extracellular leaflet.

### Secreted Neu3 disrupts neuronal network integration in primary neurons

Our lab has recently demonstrated that desialylation of primary neurons in culture by the highly promiscuous *Arthrobacter ureafaciens* (Au) sialidase results in decreased cell surface sialic acids and neuronal network integration.^21^ We hypothesized that Neu3 would have a similar effect. To assay this, we performed voltage imaging of primary neurons in culture using BeRST1,^36^ a membrane-localized voltage-sensitive fluorophore (**Figure 3a,b**). This technique enables simultaneous high-quality measurements of membrane potential in larger groups of neurons compared to traditional electrophysiology, enabling studies of network connectivity by comparing when multiple neurons in a given field of view fire.^36^ We have previously used this method in combination with factor analysis to quantify neuronal network connectivity as the “shared variance” of the network.^21^ In brief, covariance in the firing activity of measured neurons may reflect variation in synaptic input (factors), while unexplained variance reflects the fraction of the neuron’s activity that arises spontaneously. It follows then that the ratio of shared variance to total variance measures how much of a neuron’s activity is network-driven.

**Figure 3.**
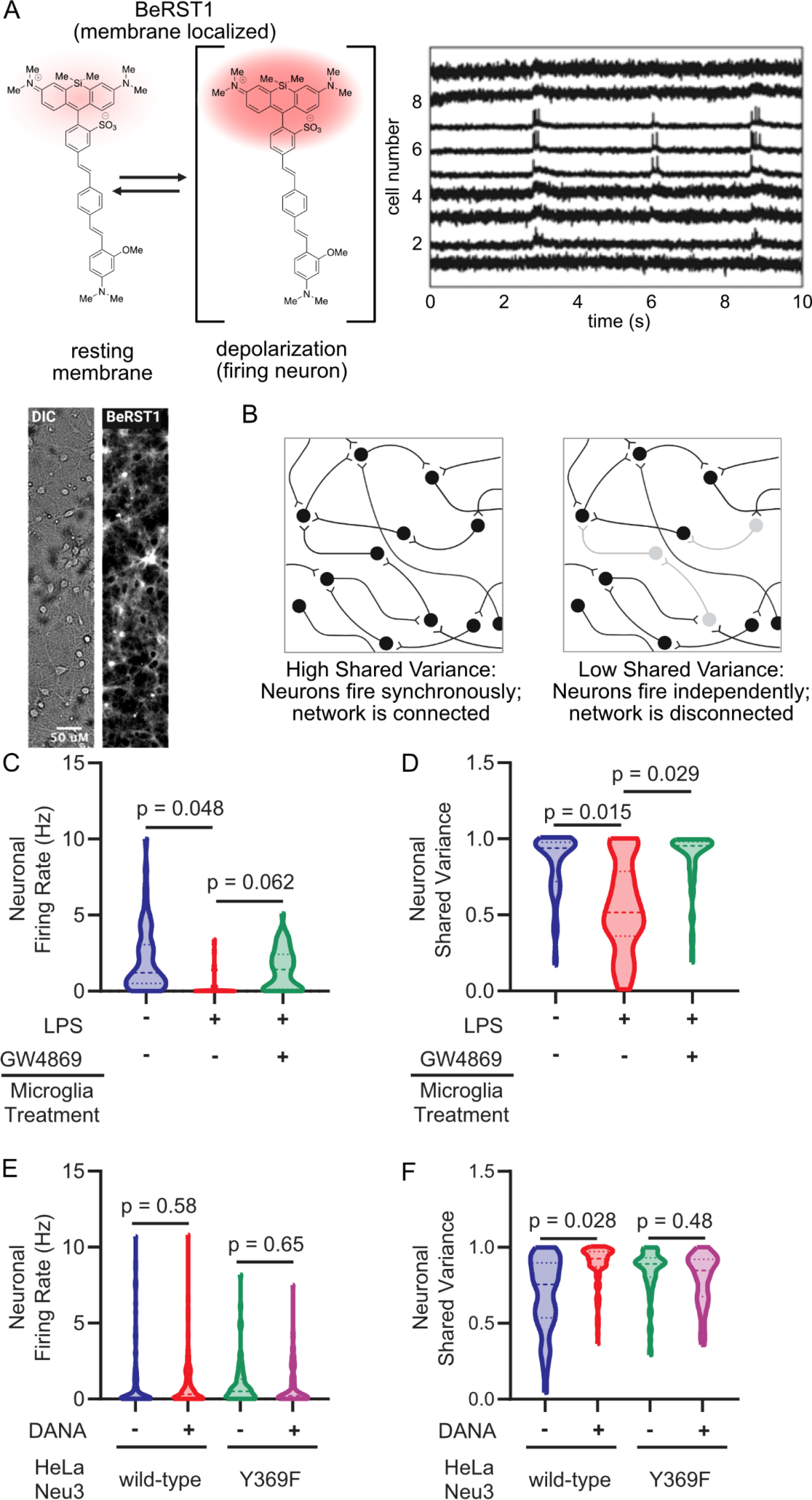
Neu3 is necessary and sufficient to disrupt neuronal network connectivity. Neurons were labeled with the voltage-sensitive dye BeRST1 and treated with extracellular-vesicle enriched media from either microglia or Neu3 over-expressing cells. Neuronal firing rates and network connectivity were analyzed by fluorescence microscopy. **(A)** BeRST1 is a membrane-localizing voltage-sensitive fluorophore that undergoes a dramatic increase in fluorescence intensity in response to changes in membrane potential, i.e. upon the depolarization of firing neurons. Representative brightfield and BeRST1 fluorescence of a single field of view and voltage traces of each neuron in a single field of view contain both subthreshold activity and spiking activity. **(B)** Network connectivity is quantitated by measuring the spike traces for individual neurons within a single field of view, and then looking at the synchronicity of firing by the metric of Shared Variance. **(C,D)** BV-2 microglia treated with or without LPS and with or without GW4869. The EVs from the conditioned media were enriched and neurons were treated with EV-enriched media, and neuronal activity was measured by voltage imaging with BeRST1. **(C)** Firing rates of neurons treated with BV-2 EV-enriched media reveal 1.7 Hz decrease in +LPS condition compared to -LPS condition (*p*=0.048). The effect is partially rescued by inhibition of EV production with GW4869 (+LPS vs. +LPS+GW4869, *p*=0.062). (**D)** Treatment with EV-enriched media results in a 29% reduction in subthreshold shared variance per neuron in -LPS vs. +LPS conditions, suggesting that neurons are no longer well-connected to the network (*p*=0.015). Effect is rescued by addition of GW4869 (+LPS vs. +LPS+GW4869, *p*=0.029). **(E,F)** As in **(C,D)**, but using conditioned media from HeLa cells overexpressing either wild-type or loss-of-function (Y369F) Neu3. **(E)** Firing rates of neurons treated with EV-enriched media of NEU3-overexpressing HeLa cells reveal no significant changes between functional NEU3 and a loss-of-function point mutant (WT: -0.26 Hz, *p*=0.58; Y369F: -0.14 Hz, *p*=0.65). **(F)** Treatment with EV-enriched HeLa media reveals 15% reduction in subthreshold shared variance between WT and Y369F mutant (*p*=0.046). Coincubation of WT EV-enriched media with DANA prevents this reduction (*p*=0.028), while coincubation of Y369F media with DANA has no significant effect (*p*=0.48). For **(C,D)**: n=4 coverslips/condition, 168 neurons total. For **(E,F)**: n=3 coverslips/condition, 331 total neurons. All hypothesis testing was performed by hierarchical permutation tests.

Using voltage imaging, we found that neurons treated with enriched EVs from activated BV-2 microglia exhibited a markedly lower firing rate compared to resting BV-2 microglia (−1.7 Hz, *p*=0.048), an effect that could be pharmacologically rescued with GW4869 (**Figure 3c**). Analyzing the data using factor analysis to quantify network connectivity, as we have previously described,^21^ we found that neuronal networks treated with activated microglial EVs experienced a substantial 29% decrease in average per-neuron shared variance compared to EVs from resting microglia (*p*=0.015). This indicates that the integration of measured neurons into a network had been significantly disrupted by treatment. This effect was rescued by pharmacological inhibition of EV production with GW4869 (*p*=0.029) (**Figure 3d**). These results indicate that activated microglial EVs are necessary and sufficient to disrupt synaptic communication.

To isolate the effects of Neu3 over other potential regulatory components of microglial secreted EVs, we overexpressed Neu3 in HeLa cells, which have been previously described to secrete Neu3 on the exterior surface of microvesicles and exosomes upon overexpression of *NEU3*.^37^ We transiently transfected HeLa cells with plasmids encoding either wild-type Neu3 or a catalytically inactive point mutant (Y369F) (**Figure S6**) and enriched EVs from the conditioned media as we did with the BV-2 conditioned media. Using periodate labeling, we observed that these EV-enriched fractions were still capable of desialylating neuronal membranes (**Figure S7**), suggesting that Neu3 is the sialidase remodeling the neuronal glycocalyces in microglial secretions. Using voltage imaging, we observed that neither wild-type or Y369F Neu3 containing HeLa-derived EV’s caused significant changes in firing rate (WT: -0.26 Hz, *p*=0.58; Y369F: -0.14 Hz, *p*=0.65), indicating that the observed decrease in firing rate of neurons treated with EV’s from activated microglia is likely due to secreted factors besides of Neu3 (**Figure 3e, S8**). However, factor analysis revealed a 15% decrease in per-neuron shared variance between WT and Y369F-treated cultures (*p*=0.046), indicating a Neu3 activity-dependent loss of connectivity (**Figure 3f**). Congruent with this, pharmacological inhibition of Neu3 with DANA abrogated the effect of wild-type Neu3 on neuronal connectivity in culture but had no significant effect in Y369F-treated cultures (WT, *p*=0.028; Y369F, *p*=0.48; **Figure 3f**). This observation demonstrates that EV-associated Neu3 activity alone is sufficient to drive changes in neuronal communication, a previously unknown function of Neu3.

## Conclusions

The prototypical sialic acid, 5-*N*-acetylneuraminic acid, was named based on the observed abundance of sialic acids on the external leaflet of neurons, particularly sialylated glycolipids known as gangliosides.^38^ Neu3 is a membrane-associated glycolipid sialidase^28^ that we speculated might play a role in regulating neuronal connectivity. The data herein present a new mechanism by which microglia regulate neuronal sialylation by secretion and transfer of Neu3 via extracellular vesicles. Moreover, we show that Neu3-mediated remodeling has a dramatic impact on the connectivity of neuronal networks, providing molecular detail for a contact-independent mechanism of neuronal pruning. These findings demonstrate a novel axis by which microglia and neurons communicate. Indeed, sialoglycans may serve as a mechanistic bridge between neuroinflammation and downstream changes in electrophysiology, which would position them as potential therapeutic targets for neurological disorders. The electrical mechanism of this rewiring, as well as other neuroinflammatory signals that lead to this effect, are exciting grounds for future research.

## Acknowledgements

This work was supported by National Institutes of Health Grant GM058867. C.S.D. was supported by a National Science Foundation Graduate Research Fellowship DGE-114747 and a Stanford Interdisciplinary Gradate Fellowship affiliated with ChEM-H. N.M.R. was funded by National Institutes of Health Grant K99GM147304. Figure illustrations were created using BioRender.com.

