## Supplemental Information for "Microglia mediate contact-independent neuronal pruning via secreted Neuraminidase-3 associated with extracellular vesicles"

### Materials and Methods.

#### Cell culture

BV2 murine microglia (a kind gift from T. Wyss-Coray) were propagated in DMEM supplemented with 10% hiFBS. Microglia were maintained at a low passage number (< 10 since obtaining initial stocks) and split at or before ~ 70% confluency to avoid runaway inflammation as caused by dead or floating BV2 cells. If cells grew overconfluent, the culture was discarded. Typically, ~ 1e6 microglia were seeded in a T75 in 25 mL split every two days. Microglia were harvested using Gibco enzyme-free dissociation buffer, incubated for 5 min at room temperature, and lifted with gentle tapping and pipetting. Microglia were diluted in an equal volume of complete media, counted with a hemocytometer, pelleted by centrifugation (300 rcf, 5 min), and resuspended in complete media for subculturing.

HeLa cells (ATCC CCL-2) were propagated in DMEM supplemented with 10% hiFBS. HeLa cells were subcultured at a confluence below 80% to avoid overgrowth. Typically, HeLa cells were split 1:5 every two to three days. HeLa cells were lifted using trypsin and pelleted by centrifugation (300 rcf, 5 min), before resuspending in fresh media.

#### Preparation of microglia-conditioned media

Low passage and subconfluent BV2 microglia were harvested using Gibco enzyme-free dissociation media, incubated for 5 min at room temperature, and lifted with gentle tapping and pipetting. Microglia were diluted in an equal volume of complete media, counted with a hemocytometer, pelleted by centrifugation (300 rcf, 5 min), and resuspended in NB++ media (PN) for subculturing.

Cultures for conditioned media were seeded at 4e4 cells per $cm^2$ at 2e5 cells per mL. In some cases, microglia were stimulated with LPS (1 µg/mL). Cells were cultured at 37 °C in 5% $CO_2$ for 18 h. The media was then harvested with careful pipetting cleared of floating cells and large debris by centrifugation (500 rcf, 5 min). The clarified media was then transferred to a clean tube. In some cases, 5-N-acetyl-2,3-dehydro-2-deoxyneuraminic acid (DANA) was added to 2 mM. An equal volume of conditioned media was then added to neuronal cultures.

#### Transfection of HeLa cells and conditioning of media

HeLa cells were transfected with plasmids encoding murine Neu3 (wt or the catalytically inactive mutant Y369F) using Transit2020 according to the manufacturer's protocol. Plasmids were custom ordered from Twist Biosciences encoding murine Neu3 (wt or Y369F) C-terminally tagged with a short linker (GSGGGSGGGGSG) followed by a 3xFLAG tag. Constructs were optimized for human codons and cloned into a pCMV vector from Twist. After 24 h, the cells were washed with OptiMEM and cultured in a low-volume of OptiMEM for an additional 24 h, at which point conditioned media was harvested and EV's were isolated.

#### Isolation and labeling of extracellular vesicles

Media was prepared as described above in 25 mL of media in a T125 cell culture flask. In some cases, bulk extracellular vesicles were isolated by concentrating media in a 100 kDa spin-filter. In other cases, exosomes were specifically isolated from conditioned media using Takara Capturem EV spin columns (Takara, 635723) according to the manufacturer's instructions.

For extracellular vesicle labeling experiments, BV-2 microglia were cultured in T25 flasks in 5 mL of Dulbecco's modified eagle medium (DMEM; Gibco) and activated overnight with LPS (1 µg/mL). Supernatant was isolated and incubated with PKH67 dye for 15 minutes at 37 °C following the PKH67GL-1KT kit (Sigma)[19]. Extracellular vesicles were then isolated as described above and resuspended in Neurobasal media (Gibco) with 1% GlutaMAX and 2% B-27 supplement. Primary hippocampal neurons were treated overnight with dyed EV preparation and imaged following kit instructions.

**GW4869 treatment of microglia**

GW4869 (#D1692, Sigma) was dissolved in DMSO to make a stock solution of 0.2 mg/mL. For inhibition of exosome generation, BV-2 microglia were treated for 3 µM GW4869 for 24 hours before 24-hour LPS treatment. Culture supernatants were collected for exosome isolation and neuronal treatment.

**Periodate labeling of cell surface sialosides**

Cells were washed with HBSS solution (Gibco) and then incubated with 1 mM sodium (meta)periodate ($NaIO_4$; Sigma) in DPBS (Sigma) for 30 minutes on ice. Cells were then washed twice with sodium acetate buffer (pH 4.7) and fixed for 10 minutes in a 1:1 acetate buffer:methanol solution on ice, then fixed for 10 min in pure methanol. Cells were then washed with sodium acetate buffer and incubated with AlexaFluor 488 hydroxylamine dye (25 uM in sodium acetate buffer; Thermo) for 1 hour at 4 Celsius. Imaging was performed on an Axio Observer Z-1 (Zeiss) equipped with a Spectra-X Light engine LED light (Lumencor), controlled with Micro-Manager. Images were acquired with a Plan-Apochromat 20x/0.8 air objective (Zeiss) paired with image capture from the ORCA-Flash4.0 sCMOS camera (sCMOS; Hamamatsu). Dye was excited with cyan LED 470/24 nm) and emission was collected after passing through the Zeiss 90 HE filter set (425/30 nm, 514/30 nm, 592/25 nm, 709/100 nm LP).

**Generation of CRISPR KO of mNeu3**

Plasmid constructs encoding Cas9 and a sgRNA were prepared in the lentiCRISPRv2 vector according to published protocols.[39] Guides were selected based on the genome-wide guides described by Bassik and coworkers.[40] Plasmid-bearing Stbl3 *E. coli* were grown in 50 mL cultures and DNA was extracted and endotoxin purified by MiraPrep.[41]

LentiCRISPRv2 plasmids were packaged in lentiviruses produced from HEK293Ts cotransfected with pGag/Pol, pRev, pTat, and pVSVG (gift from the Yi-Chang Liu and Jonathan Weismann). In brief, 1.5 µg of LentiCRISPRv2 plasmid were combined with 0.1 µg of each packaging plasmid and Lipofectamine LTX (Thermo Fisher, 15338100) in OptiMEM (Thermo Fisher, 31985062). Transfection complexes were added to HEK293Ts at 70-80% confluency in a 6 well plate in 2 mL fresh media. The media was aspirated after 12 h and bleached. After 48 h, the media was harvested and filtered through a 0.45 µm syringe filter to afford the viral supernatants.

BV2 cells were resuspended in fresh viral-containing media with polybrene (8 µg/mL). Media was changed after 24 h, and after 72 h antibiotic selection was started (2.5 µg/mL). After two weeks of selection, editing of the target gene was validated by TIDE analysis.[42]

**Cytokine release assay**

Adherent NEU3 KO and WT BV-2 microglia were plated (100,00 cells/well in a 24-well plate) in Neurobasal media (Gibco) with 1% GlutaMAX and 2% B-27 supplement one day prior to experiment. Media was treated with LPS (1 µg/mL), LPS + DANA (2 µM), or left untreated during plating. Three technical replicates were made per treatment. After 24 hours, cells were spun down at 500 rcf for 5 minutes to remove debris, and supernatant was extracted for analysis. Cytokine levels were assessed using the BD Cytometric Bead Array (CBA) Mouse Inflammation Kit. Flow cytometry was performed on a BD Accuri C6 Plus, and FlowJo software was used to gate on single cells and live cells for analysis.

µL

**Generation of endogenous tagging of mNeu3 by homology-directed recombination**

Endogenous tagging of murine Neu3 in BV2 cells was achieved using the Mendenhall-Myers system.[43] In brief, a pFETCh donor plasmid (Addgene, 63934) containing homology arms for mNeu3 (see table of gene fragments) was prepared along with PX458 plasmids containing one the only potential target cut site for mNeu3, as outlined by the target selection described by Mendenhall and Myers. Plasmids were prepared from 50 mL cultures of Stellar *E. coli* and purified by MiraPrep.[41]

Low passage BV2 microglia were transfected by magnetofection (OZ Biosciences) according to the manufacturer's protocols. After 48 h, microglia were treated with a low dose of antibiotic selection (G418, 200X). After two weeks of treatment, only the cells co-transfected with pFETCh donor and PX458 bearing the sgRNA were alive and growing well in the presence of G418 (0.25 mg/mL). The polyclonal population was grown out and the genomic DNA was isolated using a GeneJET Genome DNA Purification Kit (Thermo). PCR was performed using primers +/- 750 bp of the insertion site and compared to PCR products from wt cells. A clear 2.5 kbp band was observed in addition to a 1.5 kbp band of comparable intensity, indicating (mostly) monoallelic insertion of the transfer gene.

**Extracellular vesicles bead capture and analysis by flow cytometry**

Immunocapture beads for murine extracellular vesicles were prepared by conjugating polyclonal anti-murine CD63 (ThermoFisher, PA5-100713) to tosyl-functionalized M450 Dynabeads according to the manufacturer's protocol for antibody conjugation. After antibody conjugation, the beads were blocked with excess BSA and quenched in pH 8.5 TBS before use.

BV2-conditioned media was cleared by centrifugation (500 rcf, 5 min) and immunocaptured overnight with 2e6 anti-mCD63 beads/mL at 4 °C in a rotating Eppendorf. The beads were magnet captured and washed with cold 0.1% BSA in PBS. EV-bead complexes were stained at 2e6 beads/mL with antibody solutions for 1 h at room temperature protected from light. Antibodies were used at the following dilutions: anti-mCD9 clone KMC8 PE conjugate (ThermoFisher 12-0091-81, 1:50); anti-mCD81 clone Eat2 PE conjugate (BD Biosciences 559519, 1:50); anti-FLAG clone D6W5B AlexaFluor 647 conjugate (Cell Signaling Technologies 15009S, 1:50); rat IgG2a isotype control PE conjugate (BD Biosciences 554689, 1:50); rabbit IgG isotype control AlexaFluor 647 conjugate (ThermoFisher 51-4616-82, 1:50). Beads were magnet captured, washed, resuspended at 1e6 beads/mL and analyzed by flow cytometry on an Accuri C6 flow cytometer. Data were analyzed in FlowJo. Beads were gated on isotype-treated controls.

**Quantitative reverse transcriptase PCR analysis**

Expression levels of Neu1, Neu3, and Neu4 in murine BV2 microglia and primary murine hippocampal neurons were evaluated by quantitative RT-PCR (qPCR). Total RNA was isolated by TRIzol reagent extraction (ThermoFisher, 15596026) and Zymo RNA Clean and concentrator kits (Zymo, R1013) using manufacturer's protocols. Libraries of cDNA were generated via EcoDry premix kit (Takara, 639542) using 2 µg total RNA. Transcript levels were quantitated via qPCR using SybrGreen master mix (ThermoFisher, 4309155). Transcript levels were normalized to transcript levels of glyceraldehyde-3-phosphate dehydrogenase (GAPDH).

**Dissociated hippocampal cultures**

All animal procedures were approved by Stanford University's Administrative Panel on Laboratory Animal Care and conformed to the NIH Guide for Care and Use of Laboratory Animals and the Public Health Policy. Primary hippocampal tissue was harvested from E16.5 C57BL/6 embryonic mice (Charles River) immediately after sacrifice of the pregnant dam. The isolated hippocampi were dissociated using Papain Dissociation System (Worthington Biochemical Corporation) and trituration with fire-polished Pasteur pipettes. The dissociated cells were plated onto 12mm coverslips (Chemglass Life Sciences) pre-treated with Poly-D Lysine (PDL; 1 mg/mL, Sigma-Aldrich) at a density of 6 × 104 cells per coverslip. The cells were maintained for 24 hours in Dulbecco's modified eagle medium (DMEM; Gibco) supplemented with 4.5 g/L D-glucose, 10% FBS, 1% GlutaMAX, and

2% B-27 (Gibco), and then switched to Neurobasal media (Gibco) with 1% GlutaMAX and 2% B-27 supplement. Plates were incubated at 5% CO2, 37°C, and subsequent media changes took place every 7 days.

**Voltage imaging of neurons**

Neurons were incubated with an HBSS solution (Gibco) containing BeRST1 (1.0µM; Miller Lab, UC Berkeley) for 20 minutes at 37°C prior to imaging. Functional imaging of hippocampal cells was performed on an Axio Observer Z-1 (Zeiss) equipped with a Spectra-X Light engine LED light (Lumencor), controlled with Micro-Manager. Images were acquired with a Plan-Apochromat 20x/0.8 air objective (Zeiss) paired with image capture from the ORCA-Flash4.0 sCMOS camera (sCMOS; Hamamatsu) with 4x4 in-camera binning. We used a sampling rate of 0.5 kHz over 5 or 10 seconds in order to resolve individual action potentials. To achieve maximum sampling rate, a field of view (FOV) size of 512x100 pixels (665.6x130 µm) was used for simultaneous recording of 5-15 neurons at a time. BeRST 1 was excited with a 631 nm light (LED; 631nm, 28 nm bandpass) with an LED power of 70%. Emission from BeRST 1 was collected with a 680/10 nm bandpass emission filter after passing through a dichroic mirror (425/30 nm, 514/30 nm, 592/25 nm, 709/100 nm LP).

**Image analysis**

Analysis of voltage traces in primary neurons was performed using ImageJ and custom Python scripts. All Python scripts used to analyze data in this study are available at github.com/rishi-kulkarni/SpykeMapper. Regions of interest (ROIs) were drawn around cell bodies within a field of view and the average fluorescence over time was extracted and inputted into an Excel workbook. The fluorescence time course data was then analyzed using a custom Python script that performed subthreshold trace extraction and spike train generation. Briefly, the subthreshold activity was identified using asymmetric least squares regression and subtracted from raw time course data to generate a flattened trace containing a flat baseline and spiking activity. Spikes were identified from the flattened trace using a threshold of +6 STDEV of all cellular fluorescence values within a coverslip to generate a digitized spike train containing all-or-nothing firing data or Raster plot. Factor analysis was performed using the FactoryAnalyzer module with no rotation. The shared variance values per network were compared using Cohen’s d.

**Preparation of extracellular vesicles for mass spectrometry**

Extracellular vesicles were enriched as described above in the section Isolation and labeling of extracellular vesicles. Enriched EVs were labeled with membrane-impermeable sulfo-NHS-biotin (Thermo Fisher Scientific, 21217) per the manufacturer’s protocol. In brief, EVs in pH 8.0 PBS were labeled for 30 min at room temperature with sulfo-NHS-biotin (2 mM) and were quenched by the addition of 100 mM glycine. EVs solutions were desalted using spin columns. EVs were lysed by the subsequent addition of 10X RIPA buffer. Samples were prepared in triplicate, and triplicate control samples were treated equivalently, except sulfo-NHS-biotin was excluded. All samples were enriched using streptavidin magnetic beads (Pierce, #88817) and the DynaMag-2 Magnet (Invitrogen, #12321D) based on a protocol adopted from previous work.[44] Buffers were made fresh, including ETDA buffer (150 mM NaCl, 100 mM Tris pH 8, 0.5 mM EDTA pH 8), urea buffer (2 < urea, 10 mM Tris) and elution buffer (5% SDS in RIPA, 20 mM DTT, 2 mM biotin). 25 µL of streptavidin beads were used per sample, and beads were washed with 50 µL of cold RIPA twice before being resuspended in 25 µL cold RIPA. Beads were added to each sample, which were continuously rotated overnight at 4 C. The next day, the unbound fraction was removed from the beads and kept. Beads were washed with 100 µL RIPA, which was added to the saved unbound fraction (labeled FT, flow through) for each sample. Next, beads were washed twice with 200 µL RIPA, three times with EDTA buffer, three times with 1.5 M NaCl, three times with 0.1 M NaHCO3, and once with urea buffer. Proteins were eluted from the beads by boiling at 95 C with 60 µL elution buffer for 10 min. The elution step was repeated, for a total of two elution steps, and elutes from each step were combined for a given sample. Digestion was performed for eluted and FT proteins using a micro S-trap protocol provided by the manufacturer (Protifi).[45] For FT samples, proteins were brought to 5% SDS and reduced with 10 mM DTT for 10 minutes at 95 C. Cysteines were alkylated using 40 mM iodoacetamide for 45 minutes each at room temperature in the dark. The lysate was then acidified with phosphoric acid, brought to approximately 80-90% methanol with 100 mM TEAB in 90% methanol, and loaded onto the S-trap column. Following washing with 100 mM TEAB in 90% methanol, trypsin (Promega) was added to the S-trap at a 20:1 protein:protease ratio for 90 minutes at 47 °C. Peptides were eluted in three steps that were collected in the same tube for a given sample: 40 µL of 50 mM TEAB, 40 µL 0.2% FA in water, and 40 µL of 0.2% FA in 50% ACN, all spun at 4,000 x g for 1 minute. Eluted peptides were dried via lyophilization.

**Preparation of cell-surface labeled neurons for mass spectrometry**

Samples were prepared following published cell surface capture protocols that label cell surface proteins through sialic acids.[46,47] and lysate protein concentrations were quantitated by BCA (Pierce). Lysates were digested using a mini S-trap protocol (Protifi), which is similar to the micro S-trap protocol above, but with different volumes. Lysates were brought to 5% SDS and reduced with 5 mM DTT for 5 minutes at 95 C. Cysteines were alkylated using 25 mM iodoacetamide for 45 minutes each at room temperature in the dark. Lysates were then acidified with phosphoric acid, brought to approximately 80-90% methanol with 100 mM TEAB in 90% methanol, and loaded onto the S-trap column. Following washing with 100 mM TEAB in 90% methanol, trypsin (Promega) was added to the S-trap at a 20:1 protein:protease ratio for 90 minutes at 47 °C. Peptides were eluted in three steps that were collected in the same tube for a given sample: 80 µL of 50 mM TEAB, 80 µL 0.2% FA in water, and 80 µL of 0.2% FA in 50% ACN, all spun at 4,000 x g for 1 minute. Eluted peptides were dried via lyophilization and dried peptides were then resuspended in 100 µL 100 mM Tris, pH 8 for enrichment via streptavidin beads (Pierce, #88817) and the DynaMag-2 Magnet (Invitrogen, #12321D). Buffers were made fresh, including ETDA buffer (150 mM NaCl, 100 mM Tris pH 8, 0.5 mM EDTA pH 8). 50 µL of streptavidin beads were used per sample, and beads were washed with 200 µL 100 mM Tris three times before being resuspended in 400 µL 100 mM Tris. Beads were added to each sample (500 µL total per sample), which were continuously rotated overnight at 4 C. The next day, the unbound fraction was removed from the beads and kept. Beads were washed with 500 µL 100 mM Tris, which was added to the saved unbound fraction (labeled FT, flow through) for each sample. Next, beads were washed five times with 500 µL 100 mM Tris, five times with 500 µL EDTA buffer, five times with500 µL 1.5 M NaCl, five times with 500 µL 0.1 M NaHCO3, once with 500 µL 80% (v/v) 2-isopropanol, twice with500 µL water, and three times with 500 µL warm (60 °C) 100 mM Tris. Beads were then resuspended in 500 µL 100 mM Tris. Glycerol-free PNGaseF (New England Biolabs, # P0705L) was diluted 2-fold, and 1 µL of diluted PNGaseF was added to each set of beads. Beads with PNGaseF were incubated overnight, where PNGaseF enzymatic cleavgage release formerly N-glycosylated peptides (i.e., de-glycopeptides). Eluted de-glycopeptides were acidified with 10% FA before desalting with Strata-X reversed phase SPE cartridges (Phenomenex, #8B-S100-AAK) by conditioning the cartridge with 1 mL ACN followed by 1 mL 0.2% formic acid (FA) in water. Acidified de-glycopeptides loaded on to the cartridge, followed by a 1 mL wash with 0.2% FA in water. Peptides were eluted with 400 µL of 0.2% FA in 80% ACN and dried via lyophilization.

**Mass spectrometry proteomics LC-MS/MS**

All samples were resuspended in 0.2% formic acid in water prior to LC-MS/MS analysis, where half of the sample was injected for analysis (i.e., 5 µL or 10 µL total), and non-modified peptides and de-glycopeptides (referred to collectively as peptides in this section) were analyzed with the same LC-MS/MS method. All peptide mixtures were separated over a 25 cm EasySpray reversed phase LC column (75 µm inner diameter packed with 2 µm, 100 Å, PepMap C18 particles, Thermo Fisher Scientific). The mobile phases (A: water with 0.2% formic acid and B: acetonitrile with 0.2% formic acid) were driven and controlled by a Dionex µLtimate 3000 RPLC nano system (Thermo Fisher Scientific). An integrated loading pump was used to load peptides onto a trap column (Acclaim PepMap 100 C18, 5 um particles, 20 mm length, Thermo Fisher Scientific) at 5 µL/min, which was put in line with the analytical column 5.5 minutes into the gradient. The gradient increased from 0% to 5% B between 6 and 6.5 minutes, followed by an increase from 5% to 22% B from 6.5 to 66.5 minutes, an increase from 22% to 90% B from 66.5 to 71 minutes, isocratic flow at 90% B from 71 to 75 minutes, and a re-equilibration at 0% B for 15 minutes for a total analysis time of 90 minutes per injection. Eluted peptides were analyzed on an Orbitrap Fusion Tribrid MS system (Thermo Fisher Scientific). Precursors were ionized using an EASY-Spray ionization source (Thermo Fisher Scientific) source held at +2.2 kV compared to ground, and the column was held at 40 °C. The inlet capillary temperature was held at 275 °C. Survey scans of peptide precursors were collected in the Orbitrap from 350-1350 Th with an AGC target of 1,000,000, a maximum injection time of 50 ms, and a resolution of 60,000 at 200 m/z. Monoisotopic precursor selection was enabled for peptide isotopic distributions, precursors of z = 2-5 were selected for data-dependent MS/MS scans for 2 second of cycle time, and dynamic exclusion was set to 30 seconds with a ±10 ppm window set around the precursor monoisotope. An isolation window of 1 Th was used to select precursor ions with the quadrupole. MS/MS scans were collected using HCD at 30 normalized collision energy (nce) with an AGC target of 100,000 (200%) and a maximum injection time of 54 ms. Mass analysis was performed in the Orbitrap with a resolution of 30,000 with a first mass set at 120 Th.

**Proteomics data analysis**

Peptides from EV labeled data were searched with Morhpeus search algorithm[48] housed in the MetaMorpheus software environment (version 0.0.312)[49] using the entire mouse proteome downloaded from Uniprot[50] (reviewed, 17030 entries). Cleavage specificity was set to fully tryptic with 2 missed cleavage allowed and a minimum length of 7 residues, oxidation of methionine was set as a variable modification, and carbamidomethylation of cysteine was set as a fixed modification. Precursor and product ion tolerances were set to 5 ppm and 20 ppm, respectively. Output was set to filter at a q-value of 0.01, and FlashLFQ[51] with matching between runs was enabled. All other parameters were set as default. For quantitative comparisons, protein intensity values were log2 transformed prior to further analysis, proteins with greater than three missing values (i.e., half) per condition were removed, and missing values were imputed from a normal distribution with width 0.3 and downshift value of 1.8 (i.e., default values) using the Perseus software suite[6]. De-glycopeptides were searched with the Andromeda search engine[52] in MaxQuant.[53] Cleavage specificity was set to fully tryptic with 2 missed cleavage allowed and a minimum length of 7 residues, oxidation of methionine and deamidation of asparagine were set as a variable modification, and carbamidomethylation of cysteine was set as a fixed modification. Defaults were used for the remaining settings, including PSM and protein FDR thresholds of 0.01; and 20 ppm, 4.5 ppm, and 20 ppm for first search MS1 tolerance, main search MS1 tolerance, and MS2 product ion tolerance, respectively. The Deamidation Sites table was used for further data analysis, randomly chosen spectra were spot-checked for identification accuracy using IPSA to ensure data quality, and deamidated peptides were marked as either “True” or “False” for whether they contained the N-glycosylation sequon, N-X-S/T, where X is any residue but proline. This Deamidation Sites table file was then uploaded into Perseus and further filtered to remove 1) potential contaminants and reverse hits, 2) peptides that did not contain the N-glycosylation sequon, 3) sites with less than 0.5 localization probability, and 4) identifications that had more than three missing values. Missing values for the remaining identifications were imputed from a normal distribution with width 0.3 and downshift value of 1.8 (i.e., default values).

**Statistical analysis**

All statistical hypothesis tests were performed using either a hierarchical permutation test[54] (20,000 resamples for n=3 treatments, 70,000 for n=4, 200,000 for n>4) or a Welch’s t-test. Code is available at github.com/rishi-kulkarni/hierarch.

**DNA oligonucleotides**

| Usage | DNA Sequence |
|---|---|
| mNeu3 KO sgRNA (FWD) | CACCGGAGAGGTGCCAGATTGTGTG |

| | |
|---|---|
| mNeu3 KO sgRNA (REV) | AAACCACACAATCTGGCACCTCTCC |
| murine safe-targeting sgRNA (FWD) | CACCGGAAATCCTTACCTAAGACAA |
| murine safe-targeting sgRNA (REV) | AAACTTGTCTTAGGTAAGGATTTCC |
| mNeu3 KO PCR (FWD) | GGGCCTTCAAGATTCTGTCCATTT |
| mNeu3 KO PCR (REV) | GTGCCATGTGACTCCAAAGTCATC |
| mNeu3 KO SEQ | TTTCAATGTCCTATATTGTTTGAAAAAGAACTG |
| mNeu3 HDR sgRNA (FWD) | CACCGCGACTAAAGCCAAATCAAGA |
| mNeu3 HDR sgRNA (REV) | AAACTCTTGATTTGGCTTTAGTCGC |
| mNeu3 HDR PCR (FWD) | TCCCCGACCTGCAGCCCAGCT |
| mNeu3 HDR PCR (REV) | TGGAGAGGACTTTCCAAG |
| GAPDH qPCR (FWD) | CCCATCACCATCTTCCAGGAGC |
| GAPDH qPCR (REV) | CCAGTGAGCTTCCCGTTCAGC |
| mNEU1 qPCR (FWD) | TTCATCGCCATGAGGAGGTCCA |
| mNEU1 qPCR (REV) | AAAGGGAATGCCGCTCACTCCA |
| mNEU3 qPCR (FWD) | CTCAGTCAGAGATGAGGATGCT |
| mNEU3 qPCR (REV) | GTGAGACATAGTAGGCATAGGC |
| mNEU4 qPCR (FWD) | AGGAGAACGGTGCTCTTCCAGA |
| mNEU4 qPCR (REV) | GTTCTTGCCAGTGGCGATTTGC |

**DNA gene sequences and gBlocks**

| **Gene** | **DNA Sequence** |
|---|---|
| NEU3 HOM1 | TCCCCGACCTGCAGCCCAGCTCTACACTCGGGAAGGCTGATCATCCCCGCCTATGCCTACTATGTCTCACGTTGGTTTCTCTGCTTTGCGTGTTCAGTCAAGCCCCATTCCCTGATGATCTACAGTGATGACTTTGGAGTCACATGGCACCATGGCAAGTTCATTGAGCCCCAGGTGACAGGGGAGTGCCAAGTGGCCGAAGTGGCTGGGACGGCTGGTAACCCTGTGCTCTACTGCAGTGCCCGAACACCAAGCCGATTTCGAGCAGAGGCTTTTAGTACTGATAGTGGTGGCTGCTTTCAGAAGCCAACCCTGAACCCACAACTCCATGAGCCTCGAACCGGCTGCCAAGGTAGTGTAGTGAGCTTCCGGCCTTTGAAGATGCCAAATACCTATCAAGACTCAATTGGCAAAGGTGCTCCCGCTACTCAGAAGTGCCCTCTGCTGGACAGTCCTCTGGAGGTGGAGAAAGGAGCTGAAACACCATCAGCAACATGGCTCTTGTACTCACATCCAACTAGCAAGAGGAAGAGGATTAACCTAGGCATCTACTACAACCGGAACCCCTTGGAGGTGAACTGCTGGTCCCGCCCGTGGATCTTGAACCGTGGGCCCAGTGGCTACTCTGATCTGGCTGTTGTGGAAGAACAGGACTTGGTGGCGTGTTTGTTTGAGTGTGGGGAGAAGAATGAGTATGAGCGGATTGACTTCTGTCTGTTTTCAGACCATGAGGTCCTGAGCTGTGAAGACTGTACCAGCCCTAGTAGCGACGGGAGCGGAGGAGGTTCCGG |
| NEU3 HOM2 | AGTTCTTCTGATTCGAACATCGATGAGTGAGGCCCAGCTTCCCACAGAAAGGAATGGCAGCTACAGCCAGGGTAACAGAGGTCTCTGATGTCTAGAGAAAACTCTAAAAACTAATAATCTGCTCCTTGAATTTTTTCACTTTTCCCTTCAATGAGCATGGTGAAAATTGTGCCATATCTTACATAACGAGGCTCTTGAACTGGGAGTTTGAATCTCTTCTCTTCCCATTAAAAGGAGAGGCCATGTGCTCGCTTCGCGTTCGACAAAGCCTGGATTCTGATCTTGAGTGGAAGCCACAGGCTTGTCTTTTCCAATGGTTCACTGCTCACCTGAGTATTAGGTGATGTGTAGGTGCCTTGGCCAGAAGAAAGATCTGTGTTGTTGTATTTTTTTAAATTTATTTATTTACTATATGTAAGTACACTGCAGCTGTCTTCAGACACACCAGAAGAGGGCGTCAGATCTCATTAGAGATGGTTGTGAGCCACCATGTGGTTGCTGGGATTTGAACTCAGGACCTTCAGAAGAGCAGTCA |

| | |
|---|---|
| | GTGCTCTTAACTACTGAGCCATCTCTCAAGCCCCGCATTGCTGTATTTTTAATAA GAAAAATGCCCTTATCCTTCCAATAATGCCTGGAGCTGTACAAATTCTCTGTCTT AGAAGACTTGAGAAAGCAGAACTGTAAGGTCAGATGCTTTCTCCAGCCTTGATG CTGTGTTCCACCTTCCCTTCCTCATCCAGAAAACAGTTACTAGGGAGAAAATGA GAAACCCATGCCAGCTGCCTTGGAAAGTCCTCTCCA |
| mNEU3 wt | ATGGAAGAGGTGCCTCCTTATAGCCTTAGTAGCACCCTGTTCCAGCAAG AAGAGCAGAGTGGAGTAACTTATAGAATCCCGGCTCTTCTCTATCTCCCCCCTA CGCATACATTTTTGGCATTTGCGGAAAAAAGGACATCCGTCCGGGATGAAGAT GCGGCGTGTCTCGTGTTGAGGAGAGGACTCATGAAAGGTCGCAGTGTTCAGTG GGGGCCACAGCGCCTCTTGATGGAAGCTACGCTTCCTGGGCACAGGACAATG AACCCGTGCCCGGTATGGGAAAAGAACACTGGAAGAGTTTACTTGTTCTTCATC TGCGTGAGGGGGCACGTCACAGAGCGGTGCCAGATAGTCTGGGGAAAGAATG CGGCGCGCCTTTGTTTTTTGTGCAGTGAAGATGCGGGATGCTCCTGGGGTGAG GTGAAGGATTTGACCGAAGAAGTCATTGGCTCTGAGGTGAAAAGATGGGCAAC CTTCGCTGTCGGACCAGGACACGGGATACAGCTGCATTCAGGCCGGCTGATCA TTCCGGCTTATGCCTACTATGTCTCCCGCTGGTTCCTTTGTTTTGCATGCAGCG TCAAACCGCACTCCCTCATGATATACTCCGACGATTTCGGGGTGACATGGCATC ACGGAAAGTTCATTGAACCTCAAGTCACCGGTGAATGCCAGGTCGCGGAGGTG GCAGGTACAGCCGGCAACCCGGTCCTTTATTGCAGCGCTCGGACCCCGTCCC GCTTTAGAGCCGAAGCCTTTAGTACAGATTCTGGCGGCTGCTTCCAAAAACCG ACTCTCAACCCTCAACTCCACGAACCTAGAACAGGTTGCCAAGGAAGCGTTGT GAGCTTCCGGCCGTTGAAGATGCCAAACACATATCAAGACTCTATCGGTAAGG GGGCGCCTGCGACGCAAAAGTGTCCACTCCTCGACAGCCCACTGGAGGTCGA GAAAGGCGCGGAAACCCCTTCCGCGACGTGGTTGCTGTATTCACATCCCACTA GCAAGAGGAAGAGAATTAACCTGGGGATTTACTACAATCGCAACCCGCTGGAG GTTAACTGCTGGAGTCGGCCGTGGATCCTTAACCGGGGTCCATCAGGCTACAG CGACCTGGCGGTAGTTGAGGAGCAGGATTTGGTGGCTTGCCTCTTTGAGTGCG GGGAGAAGAACGAATATGAGCGGATCGATTTCTGTTTGTTTTCTGATCACGAAG TATTGTCATGTGAAGATTGTACTTCCCCGTCTTCAGACGGGAGCGGAGGAGGT TCCGGtggaggtggttctggagattacaaggatgacgacgataagggcgattacaaggatgacgacgataagg gagattacaaggatgacgacgataag |
| mNEU3 Y369F | ATGGAAGAGGTGCCTCCTTATAGCCTTAGTAGCACCCTGTTCCAGCAAG AAGAGCAGAGTGGAGTAACTTATAGAATCCCGGCTCTTCTCTATCTCCCCCCTA CGCATACATTTTTGGCATTTGCGGAAAAAAGGACATCCGTCCGGGATGAAGAT GCGGCGTGTCTCGTGTTGAGGAGAGGACTCATGAAAGGTCGCAGTGTTCAGTG GGGGCCACAGCGCCTCTTGATGGAAGCTACGCTTCCTGGGCACAGGACAATG AACCCGTGCCCGGTATGGGAAAAGAACACTGGAAGAGTTTACTTGTTCTTCATC TGCGTGAGGGGGCACGTCACAGAGCGGTGCCAGATAGTCTGGGGAAAGAATG CGGCGCGCCTTTGTTTTTTGTGCAGTGAAGATGCGGGATGCTCCTGGGGTGAG GTGAAGGATTTGACCGAAGAAGTCATTGGCTCTGAGGTGAAAAGATGGGCAAC CTTCGCTGTCGGACCAGGACACGGGATACAGCTGCATTCAGGCCGGCTGATCA TTCCGGCTTATGCCTACTATGTCTCCCGCTGGTTCCTTTGTTTTGCATGCAGCG TCAAACCGCACTCCCTCATGATATACTCCGACGATTTCGGGGTGACATGGCATC ACGGAAAGTTCATTGAACCTCAAGTCACCGGTGAATGCCAGGTCGCGGAGGTG GCAGGTACAGCCGGCAACCCGGTCCTTTATTGCAGCGCTCGGACCCCGTCCC GCTTTAGAGCCGAAGCCTTTAGTACAGATTCTGGCGGCTGCTTCCAAAAACCG ACTCTCAACCCTCAACTCCACGAACCTAGAACAGGTTGCCAAGGAAGCGTTGT GAGCTTCCGGCCGTTGAAGATGCCAAACACATATCAAGACTCTATCGGTAAGG GGGCGCCTGCGACGCAAAAGTGTCCACTCCTCGACAGCCCACTGGAGGTCGA |

| | |
|---|---|
| | GAAAGGCGCGGAAACCCCTTCCGCGACGTGGTTGCTGTATTCACATCCCACTA<br>GCAAGAGGAAGAGAATTAACCTGGGGATTTACTACAATCGCAACCCGCTGGAG<br>GTTAACTGCTGGAGTCGGCCGTGGATCCTTAACCGGGGTCCATCAGGCTTTAG<br>CGACCTGGCGGTAGTTGAGGAGCAGGATTTGGTGGCTTGCCTCTTTGAGTGCG<br>GGGAGAAGAACGAATATGAGCGGATCGATTTCTGTTTGTTTTCTGATCACGAAG<br>TATTGTCATGTGAAGATTGTACTTCCCCGTCTTCAGACGGGAGCGGAGGAGGT<br>TCCGGtggaggtggttctggagattacaaggatgacgacgataagggcgattacaaggatgacgacgataagg<br>gagattacaaggatgacgacgataag |

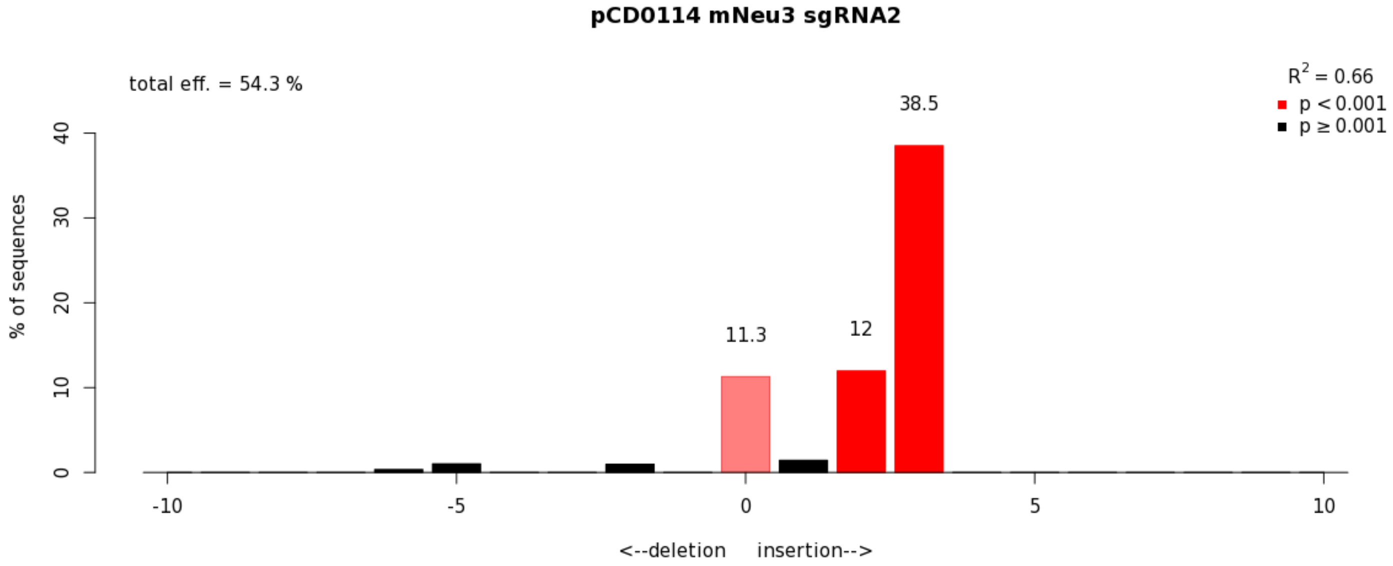


**Figure S1. Tracking of Indels by Decomposition (TIDE) analysis of *NEU3* KO BV-2 microglia.** To confirm knock-out of the *NEU3* gene in murine BV-2 microglia, CRISPR-Cas9 edited cells were subjected to two weeks of selection before genomic DNA was harvested and the cut region was amplified and sequenced. TIDE [ref] deconvolutes the possible changes from CRISPR-based editing to provide gene-level editing quantiation.

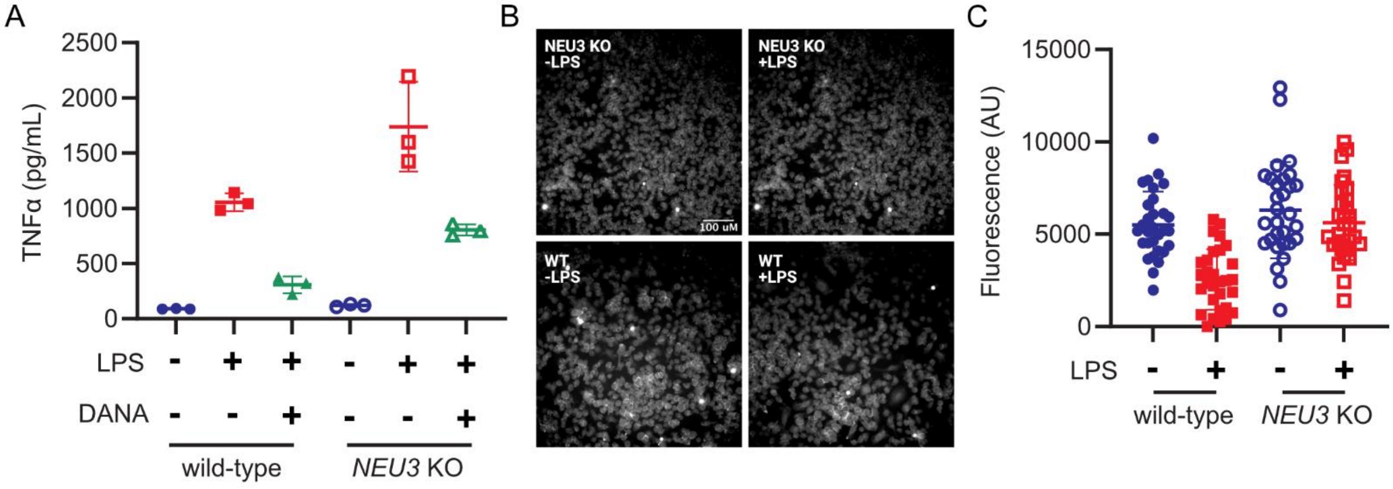


**Figure S2. Neu3 desialylates microglia in cis but is not necessary for inflammatory activity. (A,B)** Wild-type or *NEU3* knock-out (*NEU3* KO) BV-2 microglia were stimulated with LPS for 24 h, with or without deoxy-2,3-anhydroneuraminic acid (DANA). **(A)** Quantification of TNF-α secretion by LPS-treated WT and NEU3 KO microglia by flow cytometry reveals that NEU3 KO microglia are capable of activating in response to LPS (WT: -LPS vs. +LPS, $p$=0.002; -LPS vs. +LPS+DANA, $p$=0.039; +LPS vs. +LPS+DANA, $p$=0.0003) (NEU3 KO: -LPS vs. +LPS, $p$=0.02; -LPS vs. +LPS+DANA, $p$=0.001; +LPS vs. +LPS+DANA, $p$=0.055). n=3 wells/condition. Hypothesis tests performed with Welch's $t$-test. **(B, C)** Periodate labeling of activated vs. resting WT and NEU3 KO microglia (WT, $p$=0.0383; NEU3 KO, $p$=0.5073). n=3 wells/condition, 10 microglia/well. All hypothesis tests performed with hierarchical permutation test.

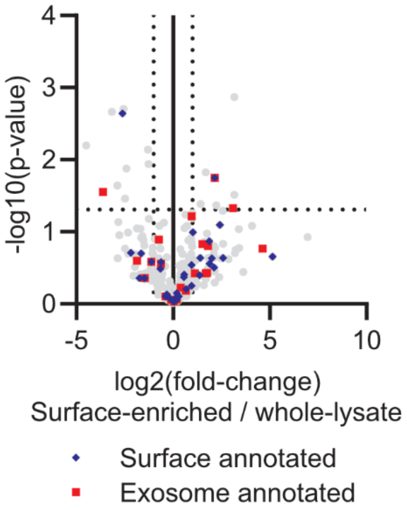


**Figure S3. Surface or whole-lysate proteomics of enriched extracellular vesicles does not identify Neu3.** Extracellular vesicles from BV-2 microglia were enriched and either lysed (whole-lysate) or subjected to cell surface biotinylation using sulfo-NHS-biotin, followed by lysis and biotin/streptavidin enrichment as described in the materials and methods. Data were quantitated by label-free quantitation. Known surface proteins (blue diamonds) and exosomal proteins (red squares), based on Uniprot GOCC annotation, are labeled.

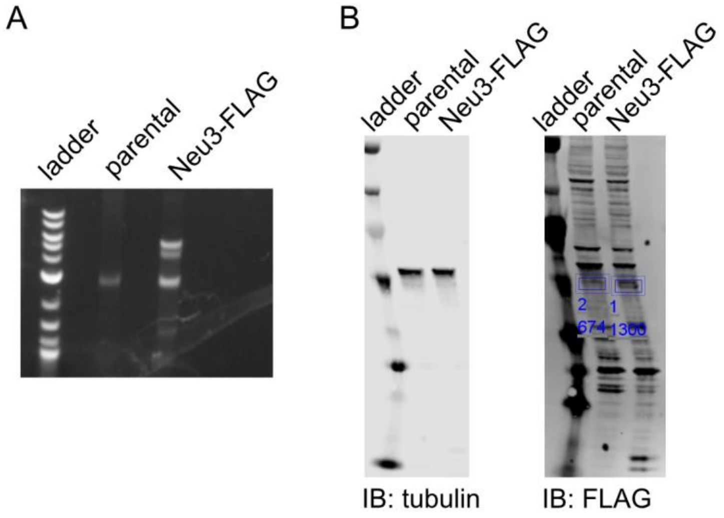


**Figure S4. Homology-directed recombination (HDR) produces endogenous Neu3 with a 3xFLAG tag. (A,B)** BV-2 microglia were co-transfected with a CRISPR-Cas9 based cutting plasmid and a donor plasmid for HDR to insert the coding sequence for a 3xFLAG tag at the C-terminus of the endogenous *NEU3* gene. (A) PCR of the NEU3 gene shows increased amplicon size in a polyclonal population of transfectants, congruent with tag incorporation. **(B)** Western-blot of whole-cell lysates from parental BV-2 or HDR-transfectants shows a noticeable and quantifiable band in the transfectants at the expected molecular weight of mNeu3.

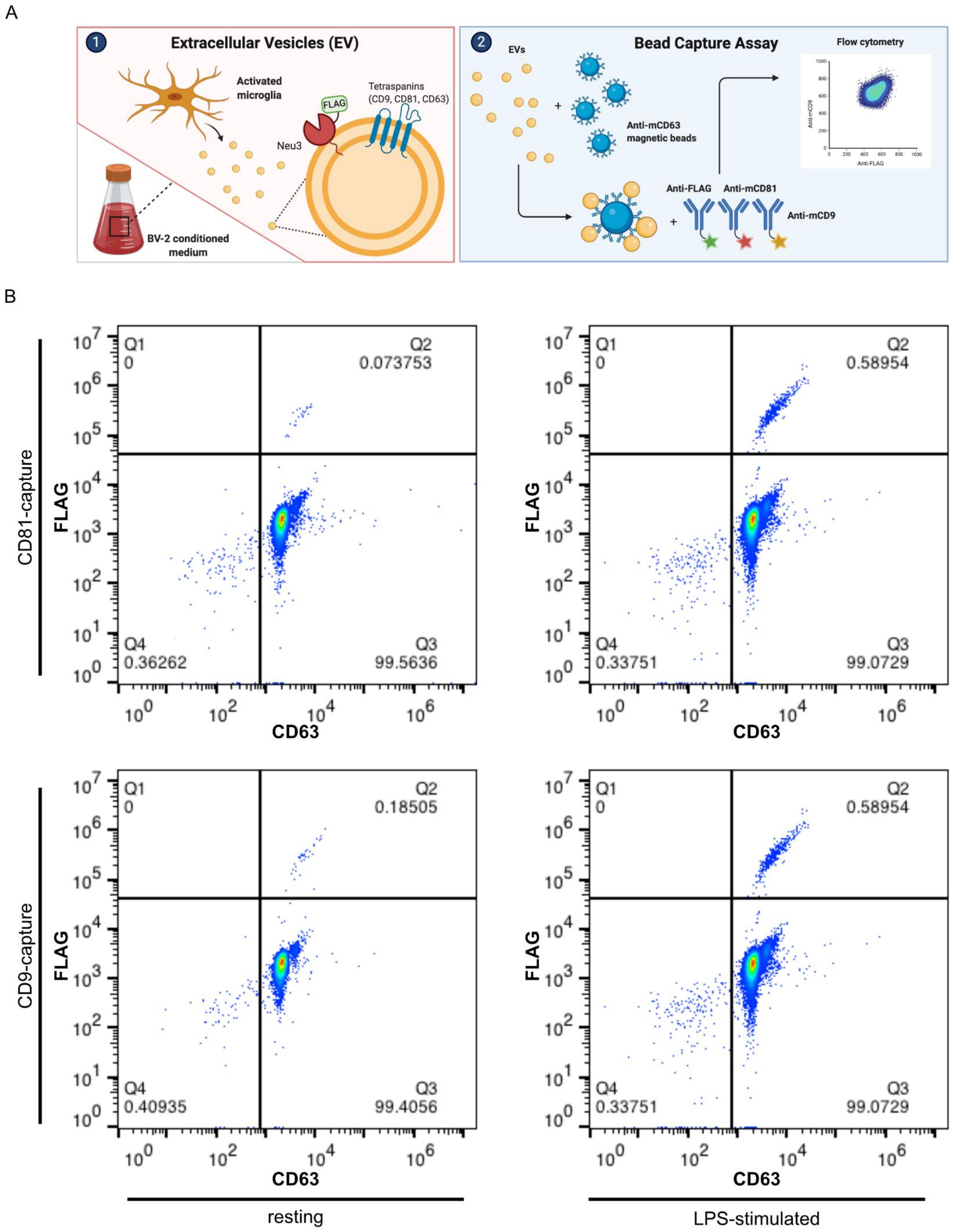
A
1
Extracellular Vesicles (EV)
Activated microglia
FLAG
Neu3
Tetraspanins (CD9, CD81, CD63)
BV-2 conditioned medium
2
Bead Capture Assay
EVs
Anti-mCD63 magnetic beads
Flow cytometry
Anti-mCD9
Anti-FLAG
Anti-FLAG
Anti-mCD81
Anti-mCD9
B
CD81-capture
CD9-capture
FLAG
CD63
Q1 0
Q2 0.073753
Q4 0.36262
Q3 99.5636
Q1 0
Q2 0.58954
Q4 0.33751
Q3 99.0729
Q1 0
Q2 0.18505
Q4 0.40935
Q3 99.4056
Q1 0
Q2 0.58954
Q4 0.33751
Q3 99.0729
resting
LPS-stimulated

**Figure S5. LPS-stimulation increases a population of Neu3+ extracellular vesicles in BV-2 microglia.** After exposure of BV-2 microglia with endogenously FLAG-tagged Neu3 to vehicle or LPS, EVs were captured on anti-mCD81 or anti-mCD9 coupled beads, labeled with fluorophore-coupled anti-FLAG or anti-mCD63, and analyzed by flow cytometry. **(A)** Representative experimental scheme. **(B)** Representative pseudocolor dot plots from bead-capture experiments in **Figure 2A**.

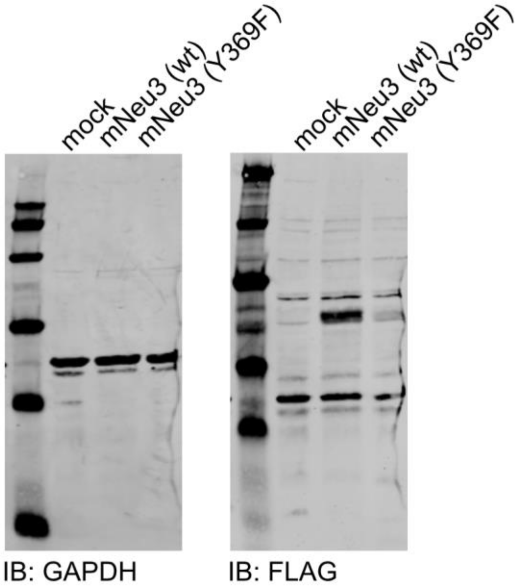


**Figure S6. HeLa cells overexpress FLAG-tagged mNeu3 upon transient transfection.** HeLa cells were transfected with plasmid encoding either wild-type mNeu3 or a loss-of-function mutant (Y396F) with a C-terminal FLAG tag. Whole cells were lysed and lysates were separated by SDS-PAGE, transferred to nitrocellulose, and probed by Western blot with IR-dye conjugated anti-FLAG or anti-GAPDH antibodies.

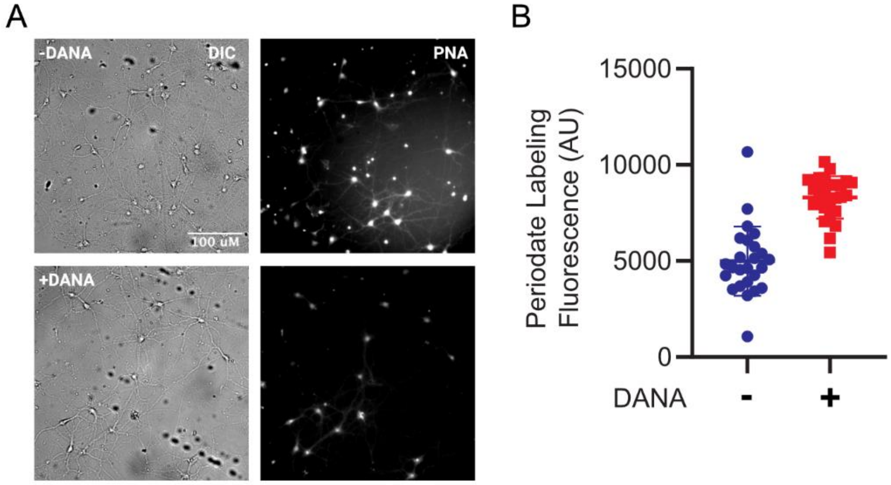


**Figure S7. Neu3 on HeLa-derived extracellular vesicles is sufficient to desialylate neurons in culture.** Primary mouse hippocampal neurons were treated with conditioned media from NEU3-overexpressing HeLa cells in the presence or absence of deoxy-2,3-anhydroneuraminic acid (DANA). Cell surface sialic acid levels were visualized by periodate labeling. Representative images **(A)** and quantification of fluorescence **(B)** reveal that Neu3 from HeLa-derived extracellular vesicles significantly reduce surface sialic acids ($p$=0.028) in a sialidase-inhibitor dependent manner. n=3 coverslips/condition, 25 cells/condition.

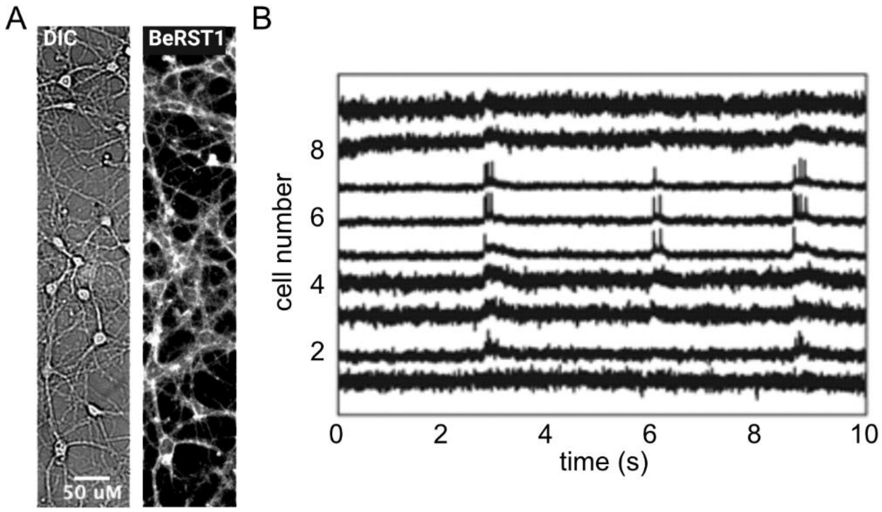


**Figure S8. Neu3 on HeLa-derived extracellular vesicles does not produce significant change in firing rate of neurons.** Representative images **(A)** and spike traces **(B)** from **Figure 3E** and **3F**.
